# Pair bonding slows epigenetic aging and alters methylation in brains of prairie voles

**DOI:** 10.1101/2020.09.25.313775

**Authors:** Lindsay L. Sailer, Amin Haghani, Joseph A. Zoller, Caesar Z. Li, Alexander G. Ophir, Steve Horvath

## Abstract

The quality of romantic relationships can be predictive of health consequences related to aging. DNA methylation-based biomarkers of aging have been developed for humans and many other mammals and could be used to assess how pair bonding impacts aging. Prairie voles (*Microtus ochrogaster*) have emerged as a model to study social attachment among adult pairs. Here we describe DNA methylation-based estimators of age for prairie voles based on novel DNA methylation data generated on highly conserved mammalian CpGs measured with a custom array. The multi-tissue epigenetic clock for voles was trained on 3 tissue sources (ear, liver, and samples of brain tissue from within the pair bonding circuit). A novel dual species human-vole clock accurately measured relative age defined as the ratio of chronological age to maximum age. According to the human-vole clock of relative age, sexually inexperienced voles exhibit accelerated epigenetic aging in brain tissue (*p* = 0.02) when compared to pair bonded animals of the same chronological age. Epigenome wide association studies identified CpGs in four genes that were strongly associated with pair bonding across the three tissue types (brain, ear, and liver): *Hnrnph1, Fancl, Fam13b*, and *Fzd1*. Further, four CpGs (near the *Bmp4* exon, *Eif4g2* 3 prime UTR, *Robo1* exon, and *Nfat5* intron) exhibited a convergent methylation change between pair bonding and aging. This study describes highly accurate DNA methylation-based estimators of age in prairie voles and provides evidence that pair bonding status modulates the methylome.

## INTRODUCTION

A significant aspect of human nature is forming social relationships between family members, friends, and romantic partners. High quality social relationships are frequently thought to be powerful predictors of long-term health, well-being, and lifespan ^1^. Marriage is perhaps the most obvious social relationship among humans, and the quality of marital relationships are known to moderate health outcomes and emotional well-being, sometimes in a sex-specific way ^2-4^. The relative risk of death, for example, is increased in non-married people compared to married people, and middle-aged males are at particular risk for several health problems, including cancer, cardiovascular disease, and respiratory diseases ^5^. In addition to cardiovascular disease, poor marital quality is related to poor health, anxiety and depression, and negative family interactions in general predict biobehavioral reactivity for anxiety, depression, and allostatic load ^6,7^. Marital dysfunction is associated with higher rates of poor diet and physical activity, which over time degrades physical health and can help explain the mechanisms by which social relationships impact health ^8^. The strength of social relationships are not only strong predictors of the risk of death, but survival and longevity are increased by strong social relationships ^9^. Thus, longevity and health across the lifespan are clearly susceptible to social support and the quality of social bonds.

Despite the persuasive evidence consistent with the idea that social relationships impact aging and lifespan, conducting empirical studies providing direct support or causal evidence for this belief is challenging. Rodent models have been of great use in studies focused on longevity and healthy aging, because they have significantly shorter lifespans by comparison to humans. However, most rodents lack the propensity to form pair bonds, a key feature that is so definitive of humans ^10^ and relatively rare among mammals in general ^11^. Prairie voles (*Microtus ochrogaster*), on the other hand, are an excellent model species for investigating the neurobiology of complex social behaviors because they form long-term socially monogamous bonds with their mates ^12,13^, and both parents exert significant and relatively equal effort to raise their young ^14,15^. In laboratory settings, prairie voles that cohabitate and mate with an opposite-sex partner for an extended period (>24 hours) exhibit a robust preference for their partner over a stranger and develop selective aggression toward unfamiliar intruders, whereas sex naïve individuals do not demonstrate these behaviors, which are consistent with a pair bond ^16^. Moreover, prairie voles that have lost a mate show depressive-like behaviors consistent with ‘grief’ and display impaired pair bond-related behaviors ^17^. The life expectancy of prairie voles in the wild fluctuates between 2 to 5 months, due to factors such as population density, season of birth, and natal social group structure ^18,19^, whereas laboratory-reared prairie voles can live up to 2-3 years. Taken together, the relatively short lifespan of prairie voles and their complex social behaviors provides a rich opportunity to examine the effects of social bonds on longevity, aging, and lifespan.

Although certain epigenetic mechanisms have recently been studied in the context of pair bonding and parental care in prairie voles ^20-24^, the impact of pair bonding on age-related epigenetic changes is an important consideration that has not received attention until now. DNA methylation (DNAm), the most studied epigenetic modification, chiefly occurs on cytosines followed by guanine residues (CpG) along the 5’→3’ direction ^25^. DNAm plays an essential role in various developmental and genomic contexts ^26-28^. Methylation of CpGs is dynamic during development and is known to regulate experience-dependent changes, such as those resulting from early-life adversity and reward-related experiences. For example, DNAm can fine-tune neuronal gene expression and social behavior outcomes in response to the type of parental care received during early postnatal development ^24^.

Growing evidence has suggested that epigenetic markers of aging based on DNA methylation data can accurately estimate chronological age for any tissue across the entire lifespan of mammals ^29-35^. These DNAm-based age estimators, also known as epigenetic clocks, target dozens to hundreds of aging-related CpG loci and apply penalized regression models to predict chronological age based on DNA methylation levels (reviewed in ^35^). By examining various mammalian tissues and cell types, a robust correlation between chronological age and DNAm age over the course of entire lifespans has been well established with the human pan tissue DNAm age estimator ^36^. Similar pan tissue clocks have been established in mice and other mammals ^33,34^.

In the present study, we used multiple tissues (brain, ear, and liver) from sex naïve and pair bonded prairie voles, of both sexes and across a wide range of ages, to develop a highly accurate pan-tissue prairie vole DNAm clock for relating DNAm age with chronological age across multiple tissues. We next assessed the degree to which the prairie vole DNAm clock is conserved, by comparing it to the human DNAm clock to assess the potential translatability of the two. We then turned our focus to determine if remaining single impacts epigenetic aging at a different rate compared to pair bonded animals (across and within specific tissues). Finally, we performed epigenome-wide association studies (EWAS) to assess the degree to which specific genes show differential epigenetic modification as a response to aging and pair bonding status.

## Results

### Data sets

We used a custom methylation array (HorvathMammalMethylChip40) to generate DNA methylation data from 141 samples from three different prairie vole tissues (brain, ear, and liver), as detailed in **Table 1**. The ages of the male and female prairie voles ranged from 0.063 to 1.11 years old. Unsupervised hierarchical clustering of the methylation data reveals that the samples cluster by tissue type (**Supplementary Figure 1**). Additionally, we used DNA methylation profiles from 850 human samples, from several tissues and with a large age range, to construct two dual species human-vole epigenetic clocks. These human data were generated on the same custom methylation array, which was designed to facilitate cross species comparisons across mammals.

**Table 1.** Description of the data. N=Total number of tissues. Number of females. Age (years): mean, minimum and maximum.

| Tissue | N | No. Female | Mean Age | Min. Age | Max. Age |
| --- | --- | --- | --- | --- | --- |
| Brain | 47 | 25 | 0.552 | 0.063 | 1.11 |
| Ear | 47 | 25 | 0.556 | 0.063 | 1.11 |
| Liver | 47 | 25 | 0.556 | 0.063 | 1.11 |

### Epigenetic clocks

To arrive at unbiased estimates of our DNA methylation-based age estimators, we performed a cross-validation study in the training data. The cross-validation study reports unbiased estimates of the age correlation R (defined as Pearson correlation between the age estimate (DNAm age)) and chronological age as well as the median absolute error. Our different clocks can be distinguished along two dimensions (species and measure of age). The pan tissue clock for voles applies to multiple tissues from voles (R = 0.94 and a median error of 0.084 years, **Figure 1A and Figure 2**). The vole pan-tissue clock is highly accurate in age estimation of the different tissue samples: R = 0.96 in brain, R = 0.95 in ear, and R = 0.91 in liver samples (**Figure 2B-D**). The human-vole pan-tissue clock for chronological age can be used to estimate the chronological age of humans and voles using the same mathematical formula. The human-vole clock exhibits a high age correlation across both species (R = 0.98, **Figure 2B**) but only moderate/weak performance when restricted to samples from voles (R = 0.57, **Figure 2C**). A better performance can be observed for the human-vole clock of *relative* age, defined as the ratio of chronological age to maximum lifespan, (R = 0.98 and R = 0.8, **Figure 2D,E**). By definition, the relative age takes values between 0 and 1 and arguably provides a biologically meaningful comparison between species with different lifespan (vole and human), which is not afforded by mere measurement of absolute age.

**Figure 1:**
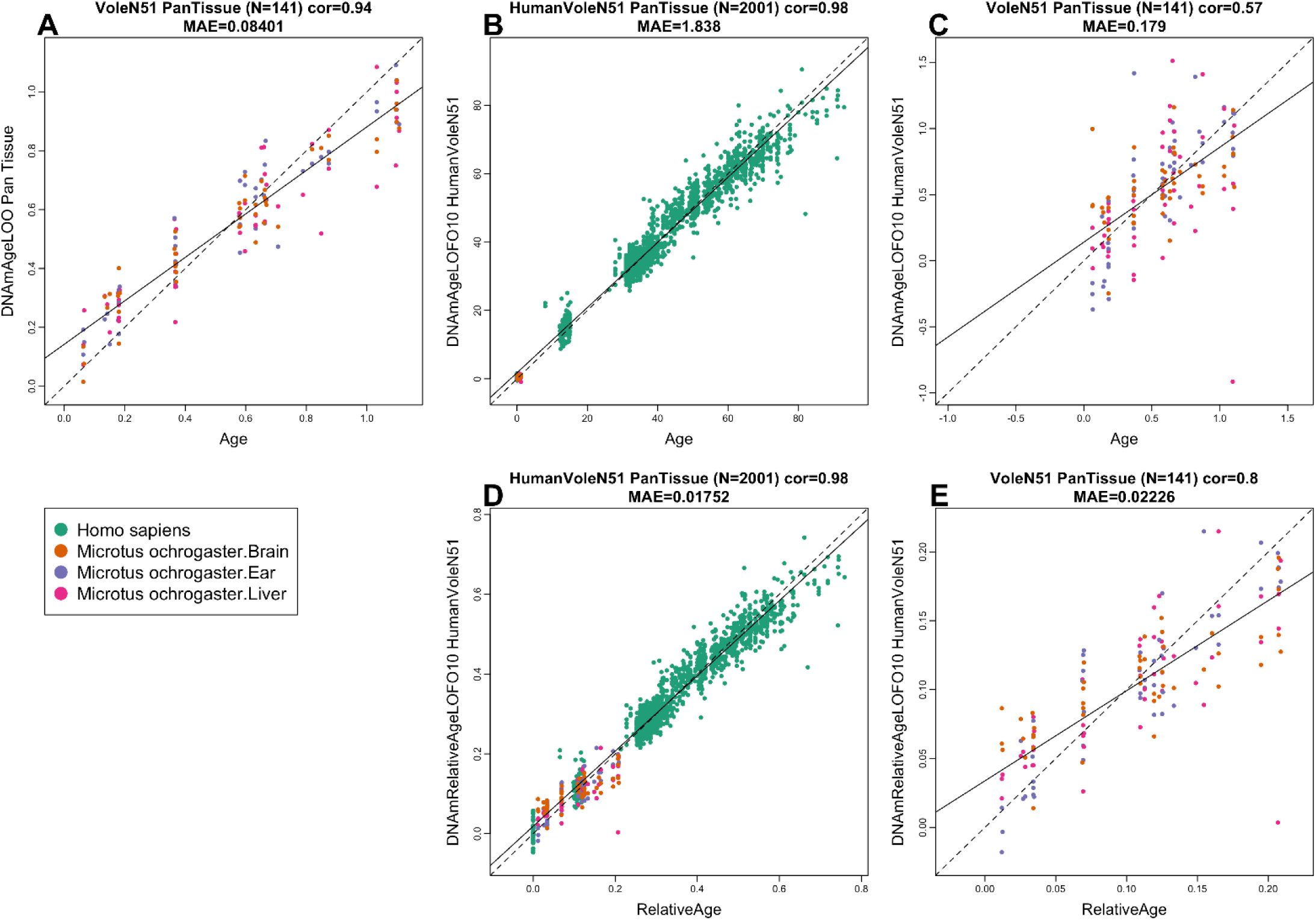
Cross-validation study of epigenetic clocks for prairie voles and humans. A) Epigenetic clock for multiple tissues from voles. Leave-one-sample-out (LOO) estimate of DNA methylation age (y-axis, in units of years) versus chronological age. B) Ten fold cross validation analysis of the human-vole clock for absolute age. C) Same clock as in panel B but restricted to voles. D) Ten fold cross validation analysis of the human-vole clock for relative age, which is the ratio of chronological age to the maximum lifespan of the respective species. E) Same clock as in panel D but restricted to voles. Dots are colored by tissue type (green=human tissue, orange=vole brain tissue, purple=vole ear tissue, pink=vole liver tissue). Each panel reports the sample size, correlation coefficient, median absolute error (MAE).

**Figure 2.**
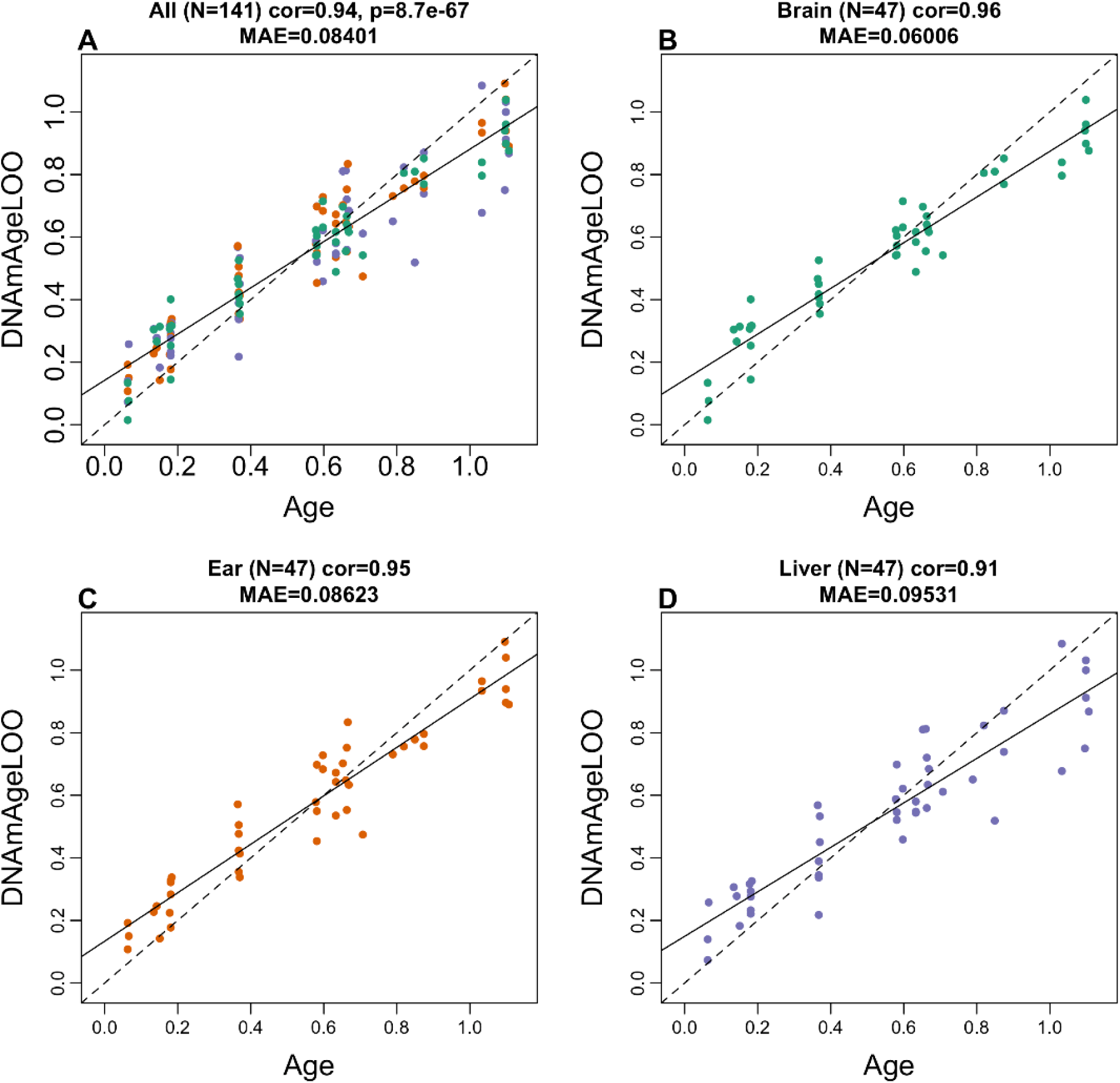
The multi-tissue epigenetic clock for prairie voles. Leave-one-sample-out (LOO) estimate of age based on DNA methylation data (x-axis) versus chronological age (in units of years) for A) all tissues, B) brain, C) ear, D) liver. Each panel reports the sample size, Pearson correlation coefficient and median absolute deviation (median error).

### Sexually experienced pair bonded voles have younger brains

We related pair bonding status (experience of being pair bonded versus sex naïve) to measures of epigenetic age acceleration in different vole tissues. Sex naïve voles were co-housed with a same-sex sibling to control for social isolation exposure that pair bonded animals received due to cohabitation with a partner; sexual experience could not be controlled because cohousing sexually experienced voles with a conspecific leads to high levels of cage-mate directed aggression. The study of pair bonding status was restricted to animals older than 0.3 years old: 20 pair bonded and 15 sex naïve. The pure pan-tissue clock for voles finds insignificant (*p* = 0.081, **Figure 3C**) but suggestive evidence that brain samples from sex naïve animals are older than those of their pair bonded counterparts (age > 0.3 years). But the human-vole clock of relative age finds nominally significant evidence (*p* = 0.02, **Figure 3D**) that older animals (> 0.3 years old) of both sexes exhibit epigenetic age acceleration in brains when compared to pair bonded animals. These results are corroborated by multivariate linear regression models that shows that pair bonded animals exhibit significantly (*p* = 0.0215) lower estimates of (cross validation-based estimates of) DNAmRelativeAge in brain samples compared to sex naïve animals even after adjusting for age and sex (**Table 2**). By contrast, the results for the pure pan-tissue vole clock were insignificant in brain samples. After increasing the age cut-off value to 0.5 years or older (> 6 months) to the pure pan-tissue clock for voles, we found suggestive evidence that sex naïve females exhibit significant age acceleration in ear tissue in comparison to pair bonded females (*p* = 0.039, data not shown). Pair bonding status was not significantly associated with epigenetic age acceleration in other tissues.

**Table 2.** Linear regression models of cross validation estimates of DNAmAge and DNAmRelativeAge. The table reports the coefficient values, standard errors, and Wald test p-values of two multivariate linear regression models. The first uses the leave-one-sample-out (LOO) estimate of DNAm age in brain samples based on the pan tissue clock for voles. The second model uses the ten-fold cross validation estimate of DNAm estimate of relative age from the human-vole relative age clock.

| Outcome: DNAmAge.LOO in brain |  |  |  |
| --- | --- | --- | --- |
|  | Coef | SE | P-value |
| Pair.bonding.statusSex naïve | 3.27E-02 | 2.33E-02 | 0.17 |
| Age | 7.10E-01 | 4.82E-02 | 1.5E-15 |
| Female | 3.44E-03 | 2.10E-02 | 0.87 |
| Outcome: Human-vole DNAmRelativeAge.LOFO10 in brain |  |  |  |
| Pair.bonding.statusSex naïve | 1.97E-02 | 8.13E-03 | 0.0215 |
| Age | 1.02E-01 | 1.68E-02 | 9.5E-7 |
| Female | 3.57E-03 | 7.32E-03 | 0.63 |

**Figure 3.**
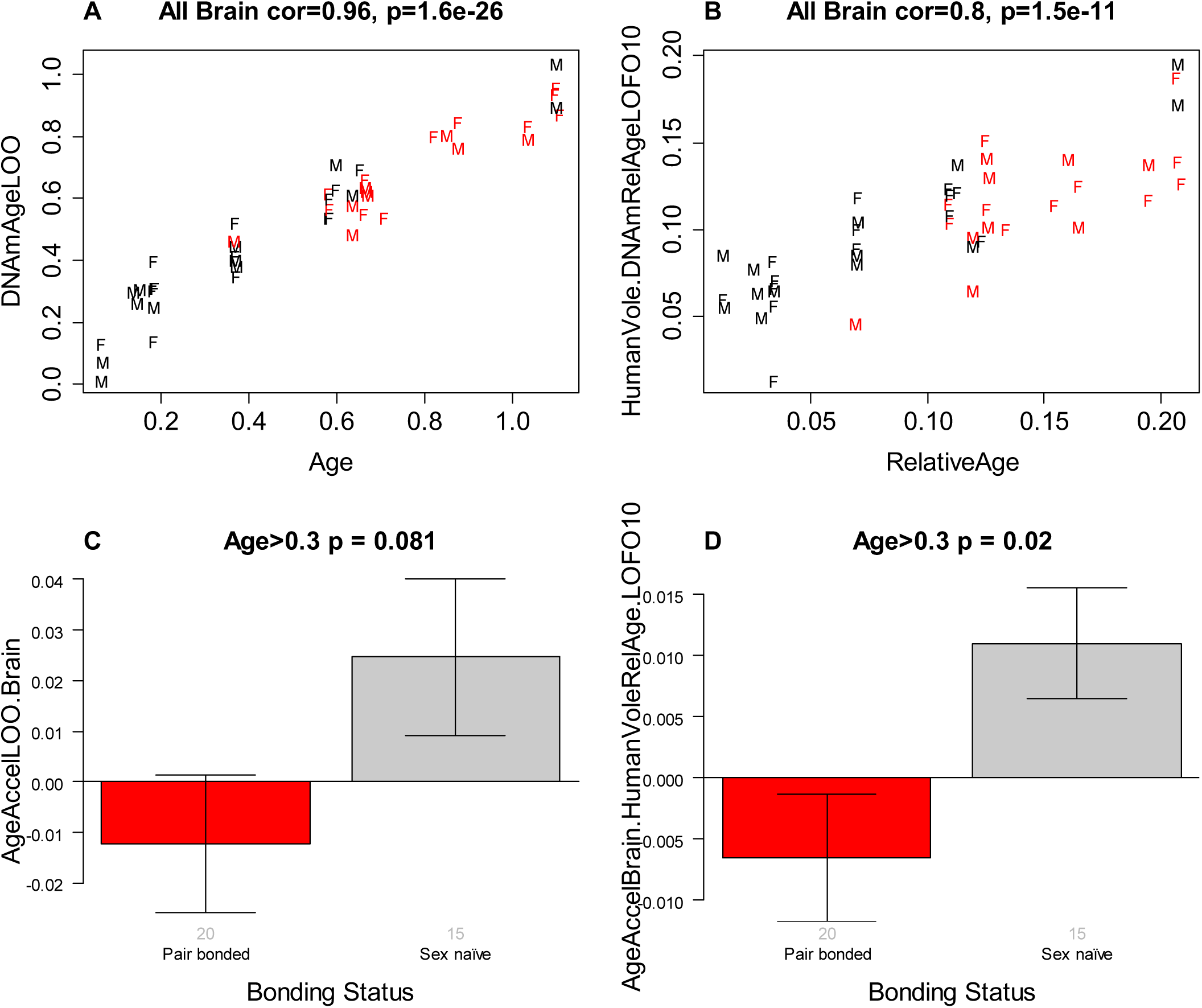
Epigenetic clock analysis of pair bonding status in brain samples. Age (axis) versus cross validation estimates of DNAmAge according to A) the pan tissue vole clock and B) the human-vole clock of relative age. Dots are colored by pair bonding status: red=pair bonded animals, black=sex naïve animals. Labels correspond to sex (F=female, M=male). C,D) Age adjusted estimates of the cross validated age estimates versus pair bonding status in animals older than 0.3 years. Bonding status versus C) epigenetic age acceleration according to the pure vole clock. D) Epigenetic age acceleration of the human vole clock for relative age. The titles of the bar plots report two-sided p-values resulting from analysis of variance. All animals younger than 0.3 were sex naïve (black color in A), which is why we restricted them from the group comparison analysis. The small grey numbers under each bar report the group sizes (n=15 sex naïve animals were older than 0.3).

### EWAS of age

In total, 33,056 probes from HorvathMammalMethylChip40 were aligned to specific loci approximate to 5,210 genes in the prairie vole genome (*Microtus ochrogaster*, MicOch1.0.100). These probes have high conservation with human and other mammalian genomes. Epigenome-wide association studies of chronological age revealed a tissue-specific DNAm change in the prairie voles (**Figure 4A**). The aging effects in one tissue seem to be poorly conserved in another tissue (**Supplementary Figure 2**). However, the poor conservation and differences in *p*-value ranges in our analyzed tissue types may reflect a limited sample size in non-blood tissues. To capture the top affected loci in all tissues, DNAm was studied at a nominal *p*-value < 10^−5^. The top DNAm changes and the proximate genic region in each tissue were as follows: brain, *Unc5a* exon (z = 8.4); ear, *En1* promoter (z = 9); and liver, *Ddx55* intron (z = −6.6). In the meta-analysis of these three tissue samples, the top DNAm changes included hypermethylation in *En1* promoter (z = 12.2), *Hoxd11* promoter (z = 10.6), *Hoxa11* exon (z = 10.4), and hypomethylation in *Tnrc6a* exon (z = −10.3). The genes with DNAm aging enriched a wide range of biological processes related to development (e.g. skeletal system development), metabolism particularly in liver (e.g. serine, glycine, threonine metabolism, and mitochondrial changes), and immune system (e.g. JUN kinase activity, cellular stress responses, and IL6 pathway) (**Supplementary Figure 3**). Moreover, aging-mediated hypermethylation seems to be labeled by H3K27ME3 marks, and regulated by polycomb repressive complex 2 (PRC2) and EED proteins. Polycomb repressive proteins regulate H3K27Me3 marks, DNA damage, and senescence states of the cells during aging ^37^. This is a conserved age-related biology among mammalian species.

**Figure 4.**
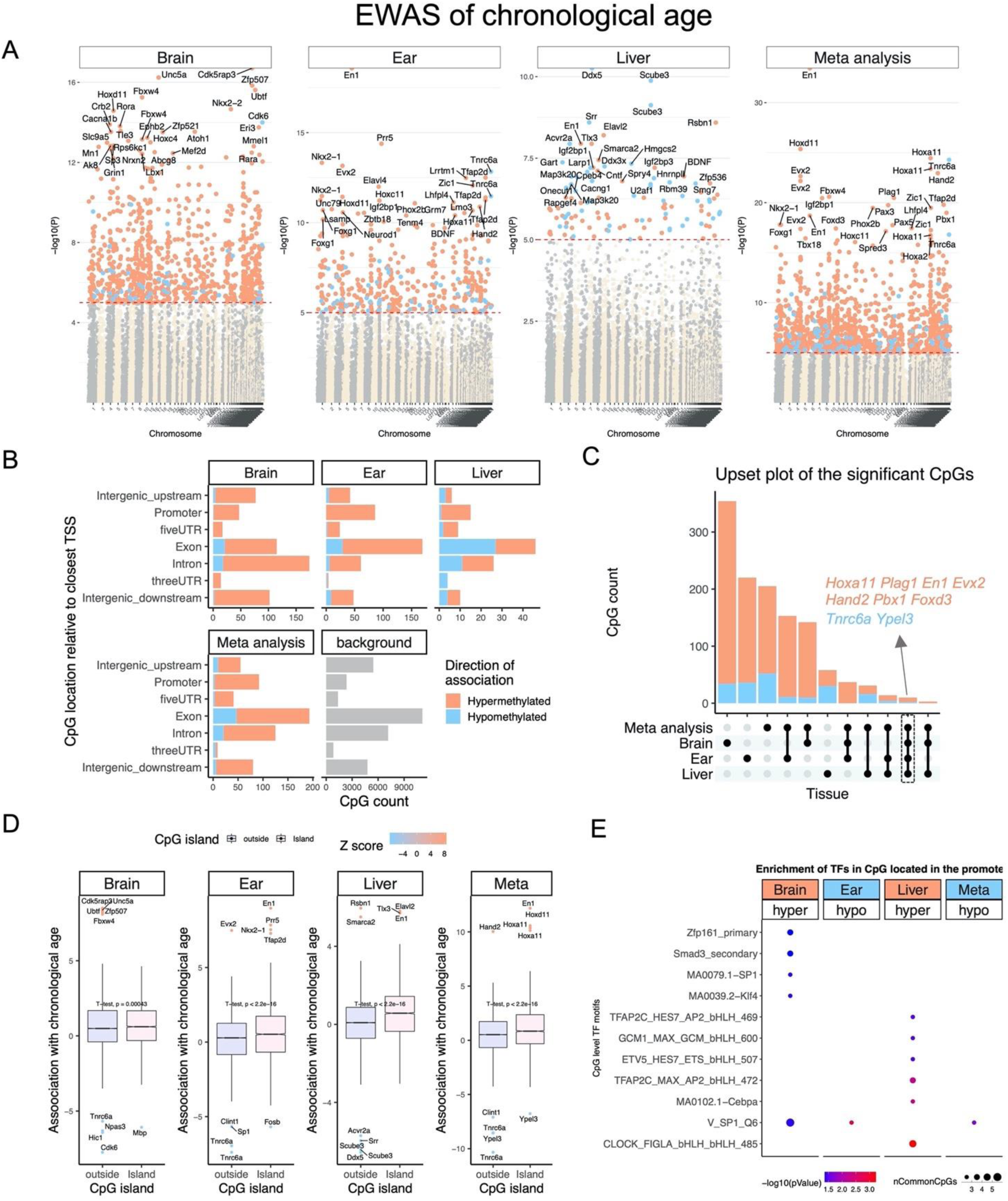
Epigenome-wide association study of age in tissues from prairie voles (*Microtus ochrogaster*). A) Manhattan plots of the EWAS of chronological age. The coordinates are estimated based on the alignment of Mammalian array probes to MicOch1.0.100 genome assembly. The direction of associations with p < ^10-5^ (red dotted line) is highlighted by red (hypermethylated) and blue (hypomethylated) colors. Top 30 CpGs was labeled by the neighboring genes. B) Location of top CpGs in each tissue relative to the closest transcriptional start site. Top CpGs were selected at p < 10-5 and further filtering based on z score of association with chronological age for up to 500 in a positive or negative direction. The number of selected CpGs: brain, 546; ear, 434; liver, 116; meta-analysis, 595. The grey color in the last panel represents the location of 33056 mammalian BeadChip array probes mapped to MicOch1.0.100 genome. C) Upset plot representing the overlap of aging associated CpGs based on meta-analysis or individual tissues. Neighboring genes of the overlapping CpGs were labeled in the figure. D) CpG islands have higher positive association with age (hypermethylation) than CpGs located outside of the islands particularly in ear and liver. E) Transcriptional motif enrichment for the top CpGs in the promoter and 5’UTR of the neighboring genes. The enrichment was tested using a hypergeometric test (Methods).

Aging-associated CpGs in different tissues were distributed in all genic and intergenic regions that can be defined relative to transcriptional start sites (**Figure 4B**). In all tissues, most of the top aging-mediated DNAm changes were hypermethylated. This pattern was more prominent in CpGs in the promoter. Further analysis using human cell type epigenetic signatures suggested that these hypermethylated CpGs are mainly located in bivalent repressors and repressed polycombs binding sites (**Supplementary Figure 4**).

Using the upset plot analysis, we identified the CpGs that showed consistent aging-mediated DNAm modifications in multiple tissues. Some of these changes included hypermethylation in *En1* promoter, and hypomethylation in *Tnrc6a* exon (**Figure 4C**). *En1* is a transcriptional factor involved in neurodevelopment ^38^. In general, CpG island showed a higher positive association with aging than CpGs located outside (**Figure 4D**). This difference was more prominent in liver and ear than brain tissues, which suggests distinct aging biology in these tissues. Transcriptional factor (TF) motif enrichment also identified distinct aging motifs between the brain and liver (**Figure 4E**). In the brain, SP1 was the top motif that showed hypermethylation with age. In contrast, SP1 was hypomethylated with age in ear samples. SP1 is a key regulator of mTORC1/P70S6K/S6 signaling pathway ^39,40^ and is involved in several aging-associated diseases including cancer ^41^, hypertension ^42^, atherosclerosis ^43^, Alzheimer’s ^44^, and Huntington diseases ^45^. In the liver, the most significant motif change was hypermethylation in CLOCK, which is involved in circadian rhythm.

### EWAS of pair bonding status and the overlap with DNAm aging

As we previously described, pair bonding status seems to alter epigenetic age acceleration in prairie vole brain tissue (**Figure 3D**). Thus, we examined pair bonded-associated specific DNAm changes, and also identified the loci that concomitantly showed age-related differences in prairie voles. Because pair bonding had a small effect size on DNAm age, the differences were studied at a nominal significance of *p* < 0.005. The total number and the most significant differentially methylated CpGs in pair bonded voles are as follows: brain, 140 CpGs, with hypomethylation of the *Fzd1* downstream region; ear, 248 CpGs, with hypermethylation of the *Galnt* upstream region; and liver, 147 CpGs, with hypomethylation in the *Tmem151b* exon (**Figure 5A**). Most of the differentially methylated CpGs showed hypermethylation by pair bonding regardless of relationship to transcriptional start sites (**Figure 5B**). The impact of pair bonding on DNAm was limited to specific loci, with minimal systematic effects on the general DNAm levels in CpG islands, or other CpGs sites (**Figure 5C**). The strongest TF motif alteration by pair bonding was hypomethylation in several immune-related motifs including ATF, CREB, FOS, and JUN in the liver (**Figure 5D**). Interestingly, pair bonding also altered some of the motifs that were jointly enriched by DNAm aging. For example, the SP1 motif was hypermethylated in the brain by both pair bonding status and aging.

**Figure 5.**
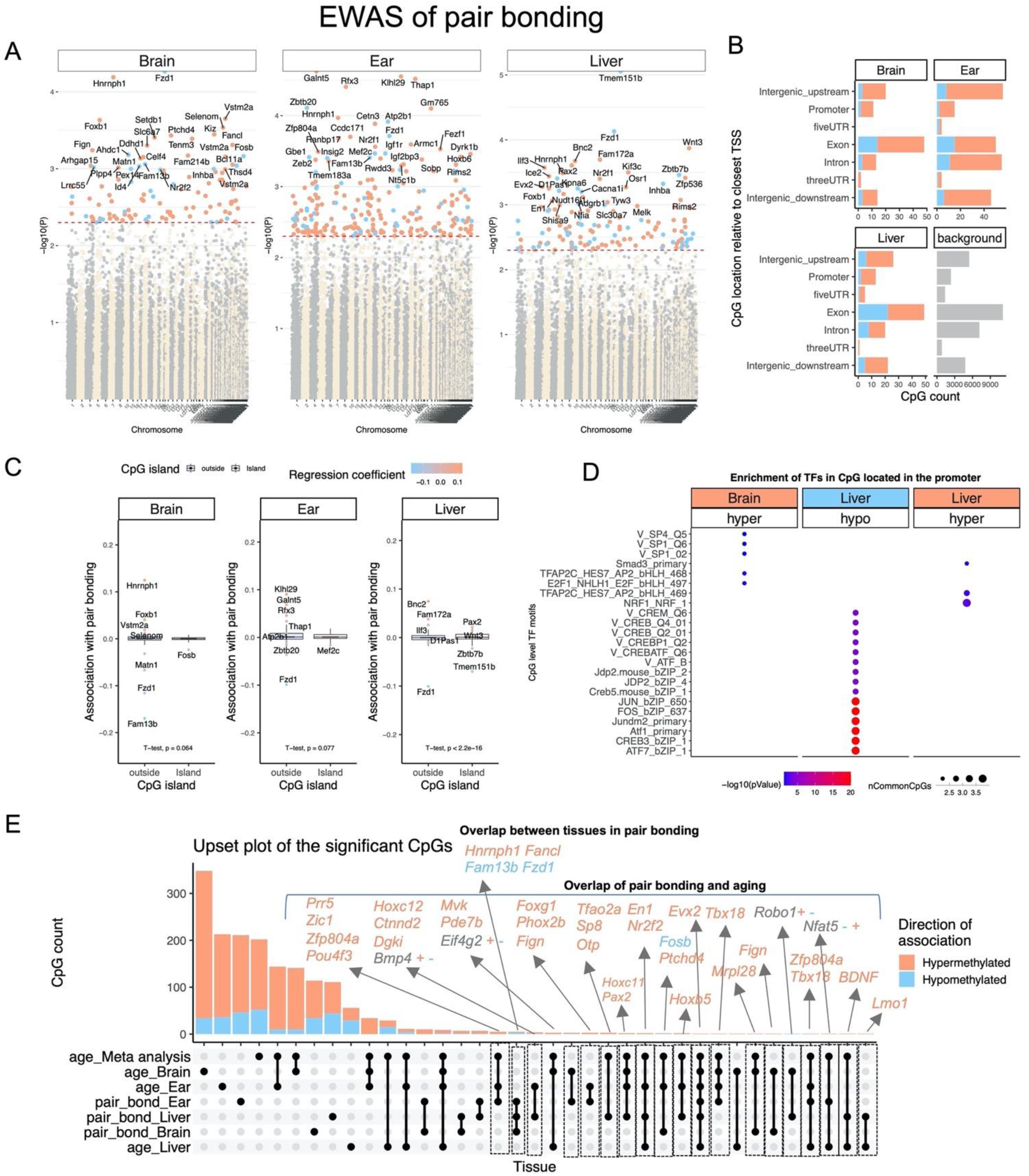
Epigenome-wide association study of pair-bonding status and the overlap with aging EWAS in tissues from prairie voles (*Microtus ochrogaster*). The association of DNAm and pair-bonding was examined by a multivariate regression model for each tissue and adjusting for chronological age as a covariate. Being single was considered as the reference for comparison with pair bonding. A) Manhattan plots of the EWAS of pair-bonding. The coordinates are estimated based on the alignment of Mammalian array probes to MicOch1.0.100 genome assembly. The direction of associations with p < 0.005 (red dotted line) is highlighted by red (hypermethylated) and blue (hypomethylated) colors in pair-bonded animals. The top 30 CpGs were labeled by the neighboring genes. B) Location of top CpGs in each tissue relative to the closest transcriptional start site. Top CpGs were selected at p < 0.005. The number of selected CpGs: brain, 140; ear, 248; liver, 147. The grey color in the last panel represents the location of 33056 mammalian BeadChip array probes mapped to the MicOch1.0.100 genome. C) Boxplot of DNAm association with pair-bonding by tissue and CpG island status of the CpGs. D) Transcriptional motif enrichment for the top CpGs in the promoter and 5’UTR of the neighboring genes (Methods). E) Upset plot representing the overlap of pair-bonding and aging-associated CpGs. Neighboring genes of the overlapping CpGs were labeled in the figure. The grey labels with + and – signs indicated CpGs with convergent association with aging and pair-bonding. The first sign (+ or -) shows the direction of association with aging, and the second sign represents the direction for pair-bonding.

Pair bonding mediated differential methylation was tissue-specific with only 4 conserved CpGs between the brain, ear, and liver (**Figure 5E**). In contrast, there were 34 conserved CpGs that were mainly hypermethylated by both pair bonding and aging in different tissues. For example, the *En1* promoter that was identified by aging-meta-analysis was also hypermethylated by pair bonding. We further identified 4 CpGs with a convergent change between pair bonding and aging. These CpGs were proximate to *Bmp4* exon, *Eif4g2* 3’UTR, *Robo1* exon, and *Nfat5* intron. The underlined loci could be the link between pair bonding- and DNAm aging-related alterations, and thus merits follow up experiments.

Gene level enrichment analysis of CpGs associated with pair bonding led to largely insignificant results (left side of **Supplementary Figure 3**). Even though these pathways overlapped with age related CpGs, the pattern seems inconclusive and might indicate a lack of sufficient gene number for GREAT enrichment analysis. Enrichment of the cell type-specific chromatin states (15 states), histone 3 marks, and DNase I hypersensitivity sites for pair bonding associated CpGs led to insignificant results (**Supplementary Figure 4**).

## Discussion

The development of prairie vole epigenetic clocks described here was based on novel DNA methylation data that were derived from 3 vole tissue types (brain, ear, and liver). We show that the pure pan-tissue vole clock accurately relates chronological age with estimated DNAm age in different prairie vole tissues. This gives us confidence that these clocks will work on new samples from other tissue types as well. A critical step toward crossing the species barrier was the use of a mammalian DNA methylation array that profiled 36 thousand probes that were highly conserved across numerous mammalian species. The prairie vole DNA methylation profiles reported here represent the most comprehensive dataset thus far of matched single base resolution methylomes across multiple tissues and ages. The two human-vole clocks estimate chronological and relative age, respectively. The dual-species clock for relative age demonstrates the feasibility of building epigenetic clocks for two species based on a single mathematical formula. This dual species clock also effectively demonstrates that epigenetic aging mechanisms are highly conserved between prairie voles and humans. The mathematical operation of generating a ratio also yields a much more biologically meaningful value of aging because it indicates the relative biological age of the organism in relation to its own species. Providing an indicator of biological age empowers the possibility to gauge potential long-term survivability with implications for reproductive fitness potential and individual mate quality.

We expect that the availability of these epigenetic clocks will provide a significant boost to the attractiveness of the prairie vole as a biological model in aging research. Prairie voles are perhaps best known for their propensity to form human-like socially monogamous pair bonds ^12,13^. The human literature has demonstrated overwhelming evidence that there are a suite of positive health and longevity benefits associated with healthy supportive marriage partnerships. We used the prairie vole as a model to understand the benefits of paired living. Our results support the human literature indicating that brains of bonded individuals age at relatively slower rates than animals that remain single. When restricting the analysis of epigenetic age acceleration to sexually mature prairie voles (> 0.3 years old), the brain tissue of sex naive animals ages faster than that of pair bonded animals according to the human-vole clock (multivariate regression model p=0.0215. Table 2). This result is largely driven by age acceleration in sex naive *males* (regression *coefficient* 0.022) as opposed to sex naïve females (regression coefficient 0.00075). These results would suggest that males are more sensitive to the preserving effects of mating partnerships. However, our data had two major limitations: 1) the two oldest sex naïve animals were males which entails that our analysis of sex naive females was underpowered, 2) overall, limited sample size: only n=7 and 8 older sex naive males and females, respectively. The human literature has found that males are more sensitive to the preserving effects of mating partnerships. Similar male-biased sensitivity among prairie voles has been reported as a result of single-gene DNA methylation associated with early life social experience on later social behavior (specifically social approach) ^24^.

EWAS analysis suggested pair bonding can alter aging biology by potentially impacting transcription factor methylation in several immune-related motifs, some immediate early genes (JUN, FosB, and c-Fos) ^46^, genes associated with neural plasticity (BDNF), and cell proliferation and apoptosis (ATF, CREB) in the brain and liver. Differential (hypo- and hyper-) methylation among immune-related motifs were particularly striking, highlighting the potential for pair bonding to impact the health status and overall quality of bonded individuals. This result might help explain the documented health benefits of supportive marriages among humans (see above). It is noteworthy that BDNF and FosB in brain were also significantly related to bonding status. BDNF is a well-known neurotrophic factor closely associated with neural plasticity, and can be highly responsive to the social environment during development and adulthood ^47,48^. *FosB* is an immediate early gene that is functionally related to the mesolimbic reward circuity and its impact on addictive behaviors ^49^. Not coincidentally, the neural structures we sampled that are part of the pair bonding circuit are also key nodes within the mesolimbic reward pathway ^50^, and it has been argued that the same circuit that modulates reward seeking behavior evolved to facilitate prosocial behaviors such as pair bonding ^51^. The functional role of *FosB* not only appears to serve as a key mediator in neural tissue associated with reward and bonding, but it also appears to relate to the advanced aging effects of bonding on the brain. Finally, the EWAS analyses identified four genes that were strongly associated with pair bonding across all three tissue types (brain, ear, and liver): *Hnrnph1, Fancl, Fam13b*, and *Fzd1*. Although their full functional significance and how they might relate to pair bonding are unclear, they could be particularly valuable targets of future study to understand how social status, such as being bonded, can impact the epigenetic motifs and functional downstream implications on gene function. Although we find promising CpGs sites that relate to aging and pair bonding status, follow up studies are needed to validate these findings and to elucidate the mechanism.

Beyond their utility, the epigenetic clocks for prairie voles reveal several salient features with regard to the biology of aging. First, the vole pan-tissue clock re-affirms the implication of the human pan-tissue clock that aging might be a coordinated biological process that is harmonized throughout the body. Second, the ability to develop human-vole pan-tissue clock for relative age attests to the high conservation of the aging process across two evolutionary distant species. This increases the likelihood, albeit does not guarantee, that conditions (such as pair bonding status) that alter the epigenetic age of prairie voles, as measured using the human-vole clock, will exert a similar effect in humans. Overall, this study provides evidence linking social monogamous life strategies with epigenetic aging in an attractive animal model.

## Materials and Methods

### Prairie vole colony

Male and female prairie voles (*Microtus ochrogaster*) were produced from laboratory-bred colonies at Cornell University, from breeding pairs that were offspring of wild caught animals captured in Champagne County, Illinois, USA. Voles are weaned and housed with littermates on postnatal day (PND) 21, and then housed with same-sex littermates after PND42-45. All animals received rodent chow (Laboratory Rodent Diet 5001, LabDiet, St. Louis, MO, USA) and water *ad libitum* and were maintained under standard laboratory conditions (14L:10D cycle, lights on at 08:00, 20 ± 2 °C) in transparent polycarbonate cages (29 × 18 × 13 cm) lined with Sani-chip bedding and provided nesting material. All experimental procedures were conducted and approved by the Institutional Animal Care and Use Committee (IACUC) of Cornell University (2013-0102) and were in accordance with the guidelines set forth by the National Institutes of Health.

### Prairie vole tissue sample collection

Ear, liver, and brain samples from the Cornell University prairie vole colony were collected from 48 male and female prairie voles at various life stages: neonatal (<1 month old), sub-adult (2-4 months old), mature adult (4-10 months old), and middle aged/old adult (>10 months old). Animals were euthanized via rapid decapitation, their tissues rapidly extracted and frozen on dry ice before being stored at −80C until further processing for genomic DNA extraction. Brains were coronally sectioned and brain regions from the pair bonding circuit (PBC) were micro-dissected and pooled for each animal. The PBC brain regions included the prefrontal cortex, nucleus accumbens, lateral septum, ventral pallidum, and medial amygdala, and ventral tegmental area ^52^. Genomic DNA was isolated and purified using the phenol-chloroform extraction and ethanol precipitation method. A total of 144 tissue samples were collected and processed for DNA methylation analysis. One animal was removed from the study due to a mismatch with the reported sex and our DNA methylation-based sex estimator.

### Human tissue samples

To build the human-vole clock, we analyzed previously generated methylation data from n=850 human tissue samples (adipose, blood, bone marrow, dermis, epidermis, heart, keratinocytes, fibroblasts, kidney, liver, lung, lymph node, muscle, pituitary, skin, spleen) from individuals whose ages ranged from 0 to 93. The tissue samples came from three sources. Tissue and organ samples from the National NeuroAIDS Tissue Consortium ^53^. Blood samples from the Cape Town Adolescent Antiretroviral Cohort study ^54^. Skin and other primary cells provided by Kenneth Raj ^55^. Ethics approval (IRB#15-001454, IRB#16-000471, IRB#18-000315, IRB#16-002028).

### DNA methylation data

We generated DNA methylation data using the custom Illumina chip “HorvathMammalMethylChip40” following a previously described procedure (Arneson, Ernst, Horvath, in preparation). The mammalian methylation array is attractive because it provides very high coverage (over thousand X) of highly conserved CpGs in mammals. Two thousand out of 38k probes were selected based on their utility for human biomarker studies: these CpGs, which were previously implemented in human Illumina Infinium arrays (EPIC, 450K) were selected due to their relevance for estimating age, blood cell counts, or the proportion of neurons in brain tissue. The remaining 35,988 probes were chosen to assess cytosine DNA methylation levels in mammalian species. The particular subset of species for each probe is provided in the chip manifest file can be found at Gene Expression Omnibus (GEO) at NCBI as platform GPL28271. The SeSaMe normalization method was used to define beta values for each probe ^56^.

### Penalized Regression models

Details on the clocks (CpGs, genome coordinates) and R software code are provided in the Supplement. Penalized regression models were created with glmnet ^57^. We investigated models produced by both “elastic net” regression (alpha=0.5). The optimal penalty parameters in all cases were determined automatically by using a 10-fold internal cross-validation (cv.glmnet) on the training set. By definition, the alpha value for the elastic net regression was set to 0.5 (midpoint between Ridge and Lasso type regression) and was not optimized for model performance.

We performed a cross-validation scheme for arriving at unbiased (or at least less biased) estimates of the accuracy of the different DNAm based age estimators. One type consisted of leaving out a single sample (LOOCV) from the regression, predicting an age for that sample, and iterating over all samples. A critical step is the transformation of chronological age (the dependent variable). While no transformation was used for the pan tissue clock for voles, we did use a log linear transformation for the dual species clock of absolute age. To introduce biological meaning into age estimates of voles and humans that have very different lifespan; as well as to overcome the inevitable skewing due to unequal distribution of data points from voles and humans across age range, relative age estimation was made using the formula: Relative age= Age/maxLifespan where the maximum lifespan for the two species was chosen from the an Age data base ^58^. Maximum age of voles and humans was set to 5.3 and 122.5 years, respectively.

### Epigenome wide association studies of age

EWAS was performed in each tissue separately using the R function “standardScreeningNumericTrait” from the “WGCNA” R package. Next the results were combined across tissues using Stouffer’s meta-analysis method.

### Transcription factor enrichment and chromatin states

The FIMO (Find Individual Motif Occurrences) program scans a set of sequences for matches of known motifs, treating each motif independently^59^. We ran TF motif (FIMO) scans of all probes on the HorvathMammalMethyl40 chip using motif models from TRANSFAC, UniPROBE, Taipale, Taipaledimer and JASPAR databases. A FIMO scan p-value of 1E-4 was chosen as cutoff (lower FIMO p-values reflect a higher probability for the local DNA sequence matching a given TF motif model). This cutoff implies that we find almost all TF motif matches that could possibly be associated with each site, resulting in an abundance of TF motif matches. For more stringent TF motif matches, a cutoff of 1e-5 could also be used. We caution the reader that our hypergeometric test enrichment analysis did not adjust for CG content. Our chromatin state analysis was conducted with eForge version 2 ^60^.

## URLs

## Acknowledgements

This work was supported by the Paul G. Allen Frontiers Group (PI Steve Horvath).

## SUPPLEMENTARY MATERIAL

**Supplementary Figure 1.**
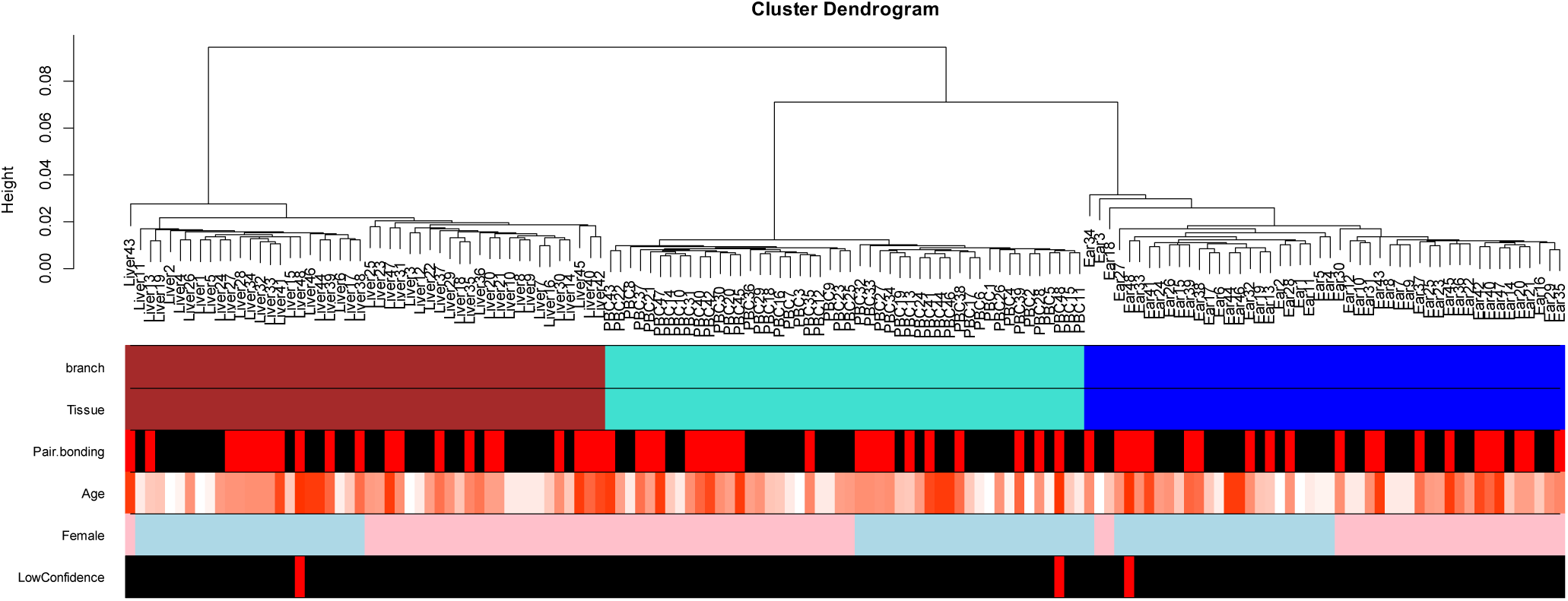
Unsupervised hierarchical clustering of tissue samples from prairie voles. Average linkage hierarchical clustering based on the interarray correlation coefficient (Pearson correlation). The first color band is based on cutting the branches at a height cut-off of 0.04. Note that the branch colors correspond to tissue (second color band): brown=liver, light blue=pooled pair bonding circuit (PBC) brains regions, dark blue=ear. The third color band visualizes pair bonding status: red=pair bonded, black=sex naïve. Pair bonding status does not seem to correspond to distinct clusters. Fourth color band visualizes age. Fifth color band visualizes sex: light blue=male, pink=female. Sixth color band visualizes 3 tissues from a single animal that were excluded from the analysis. The identity of this animal was uncertain because our DNA methylation-based sex estimator disagreed with the reported sex.

**Supplementary Figure 2.**
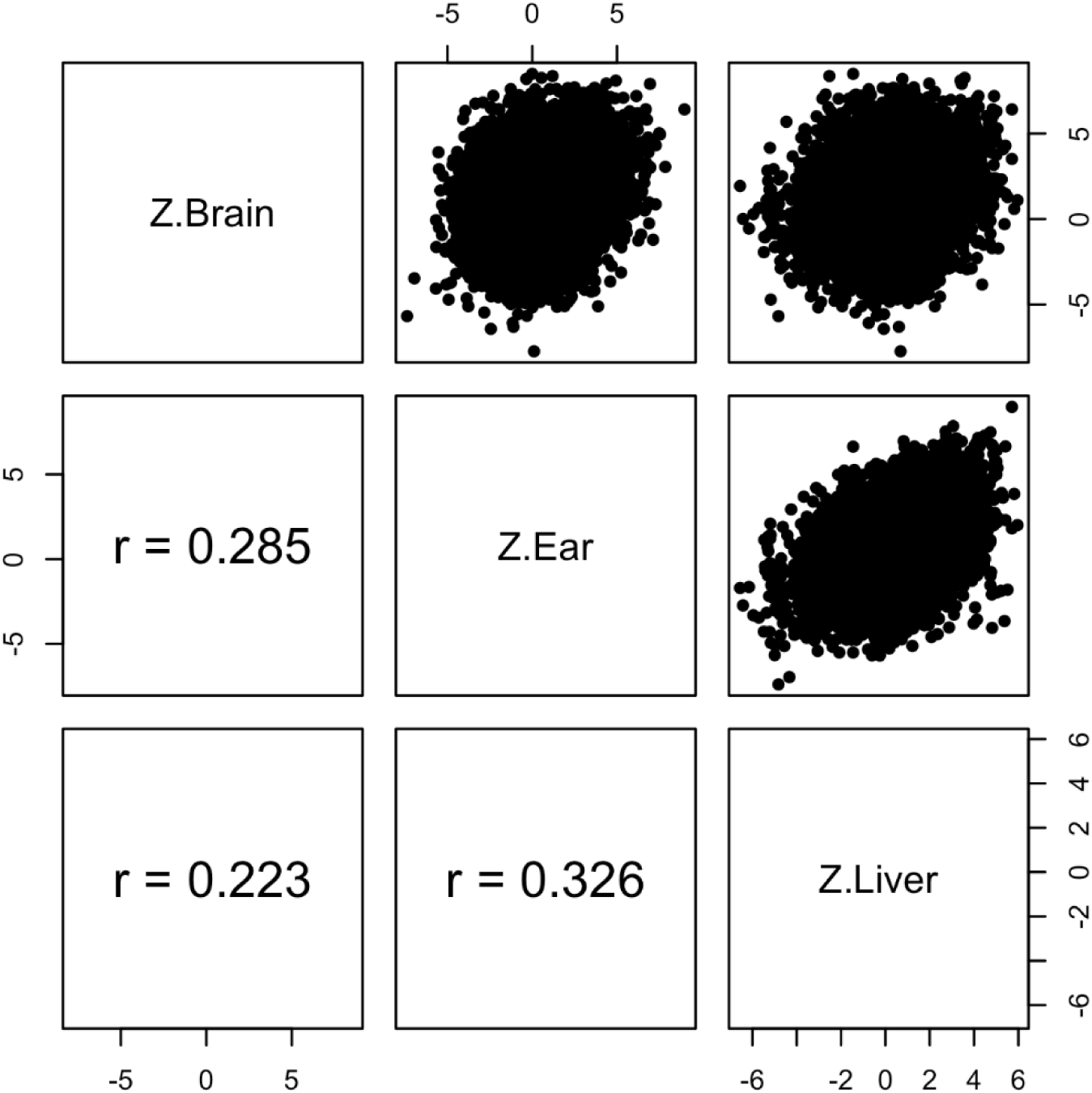
Epigenome wide association study of correlation in three different tissues. Each dot corresponds to a CpG. Z statistics for a correlation test of age in brain, ear, liver.

**Supplementary Figure 3.**
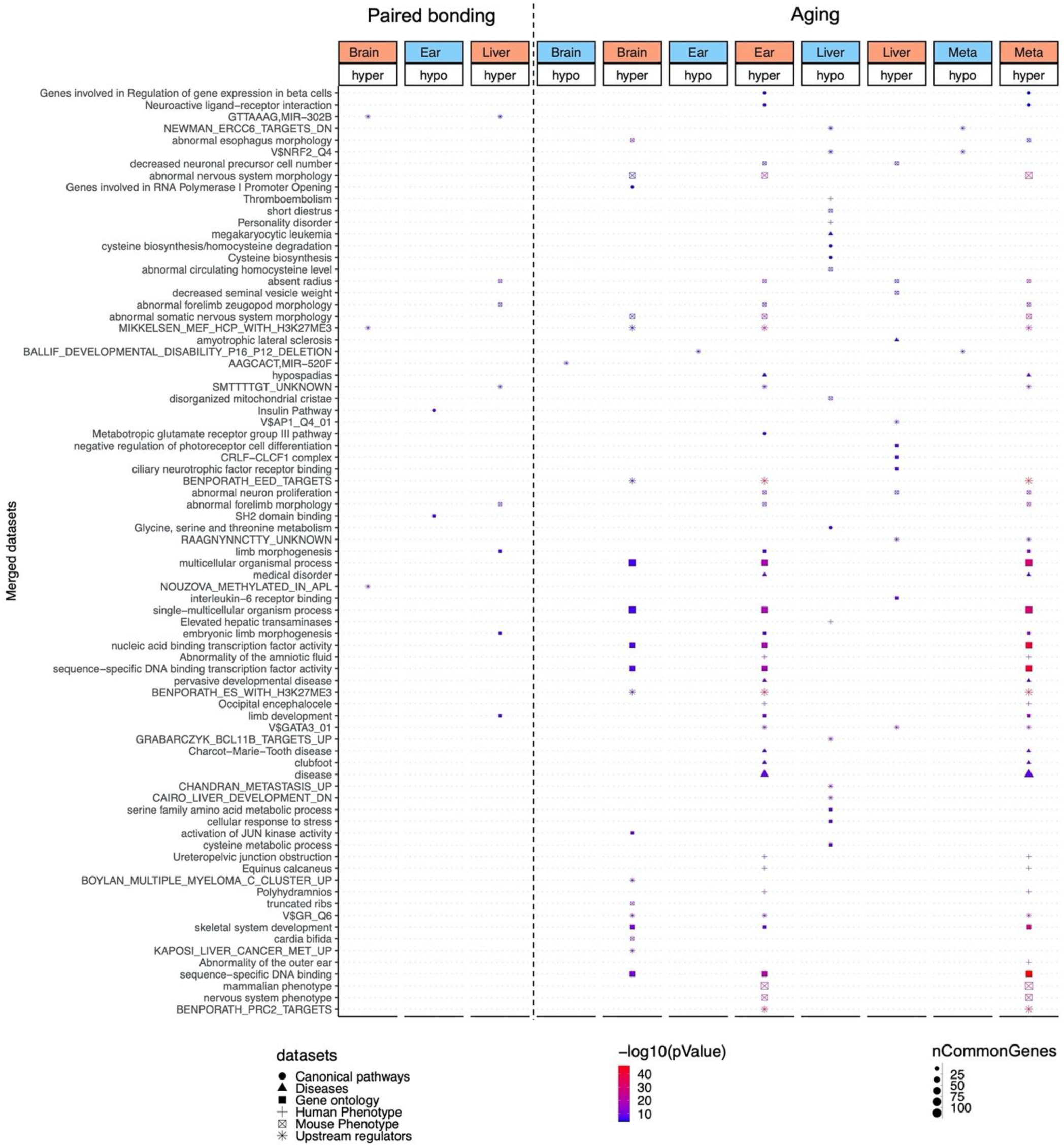
Enrichment analysis of the top CpGs with pair bonding and aging association. The analysis was done using genomic region of enrichment annotation tool ^61^. The gene level enrichment was done using GREAT analysis ^61^ and human Hg19 background. The background probes were limited to 22264 probes that were mapped to the same gene in the prairie vole genome. The top 3 enriched datasets from each category (Canonical pathways, diseases, gene ontology, human and mouse phenotypes, and upstream regulators) were selected and further filtered for significance at p < 10^−4^.

**Supplementary Figure 4.**
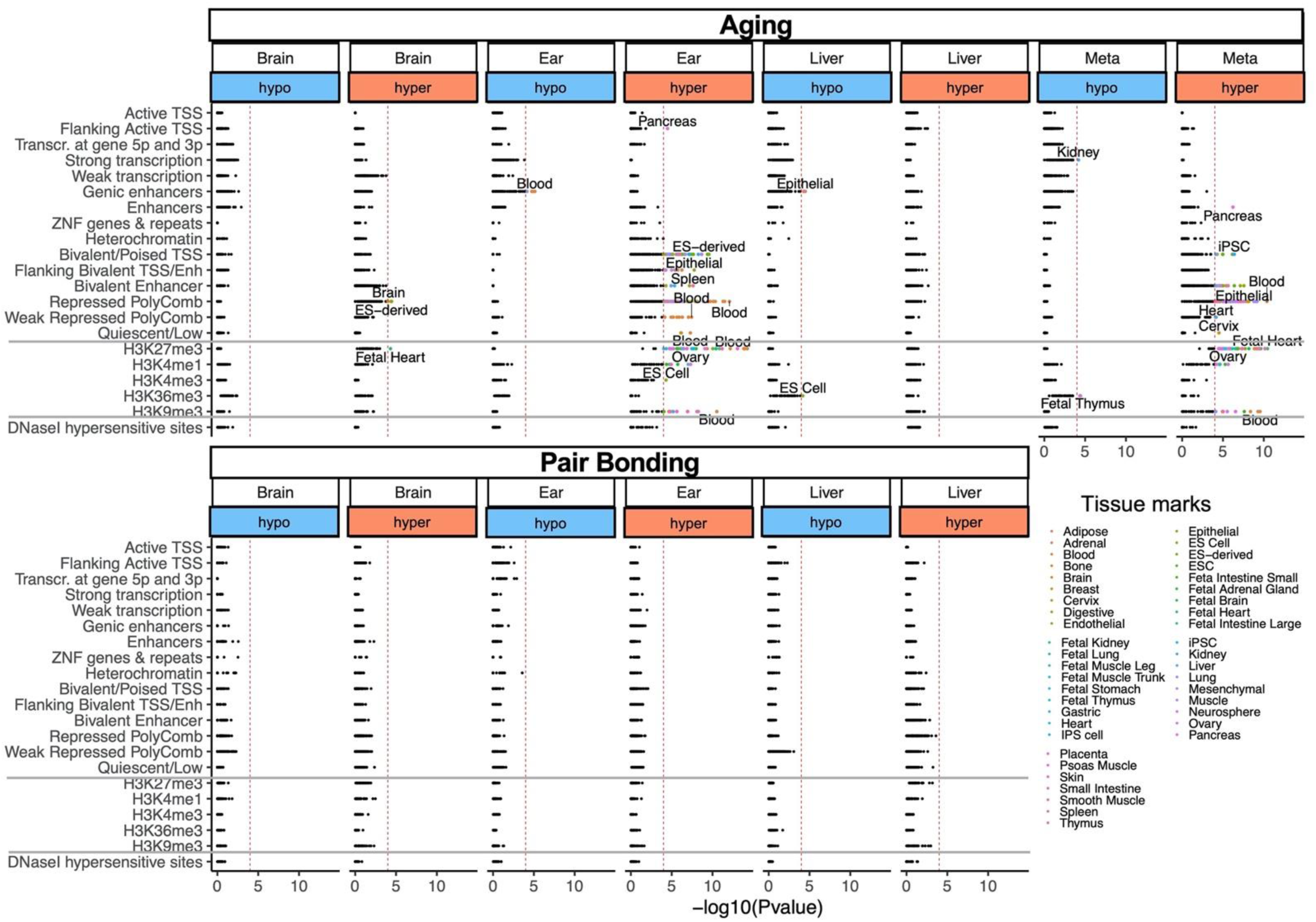
Enrichment of the cell type-specific chromatin states (15 states), histone 3 marks, and DNase I hypersensitivity sites for ageing and pair bonding-associated CpGs in prairie voles. Highlighted points indicate p < 10^−4^. The top tissue types for each significant mark are labeled. The analysis was with eForge V2.0 using *Microtus ochrogaster* genome as background.

